# Face Selective Neural Activity: Comparison Between Fixed and Free Viewing

**DOI:** 10.1101/748756

**Authors:** Carmel R. Auerbach-Asch, Oded Bein, Leon Y. Deouell

**Author notes:** Corresponding Author: Name: Carmel Ruth Auerbach-Asch.

## Abstract

Event Related Potentials (ERPs) are widely used to study category-selective EEG responses to visual stimuli, such as the face-selective N170 component. Typically, this is done by flashing stimuli abruptly at the point of static gaze fixation. While allowing for good experimental control, these paradigms ignore the dynamic role of eye-movements in natural vision. Fixation-related potentials (FRPs) obtained using simultaneous EEG and eye-tracking, overcome this limitation. Various studies have used FRPs to study processes such as lexical processing, target detection and attention allocation. The goal of this study was to compare face-sensitive activity evoked by stimulus abrupt appearance with that evoked by self-controlled gaze fixations on a stimulus. Twelve participants were studied in three experimental conditions: Free-viewing (FRPs), Cued-viewing (FRPs) and Control (ERPs). We used a multiple regression approach to disentangle overlapping activity components. Our results show that the N170 face effect (face vs. non-face) is evident for the first fixation on a stimulus, whether it follows a self-generated saccade or stimulus appearance at fixation point. Furthermore, the N170 exhibits category-specific adaptation in free viewing. The N170 face effect had similar topography across viewing conditions, but there were major differences within each stimulus category. We ascribe these differences to an overlap of the fixation-related Lambda response and the N170. We tested the plausibility of this account using dipole simulations. This study establishes the use of the N170 face effect as a signature of face detection in free-viewing experiments while highlighting the importance of accounting for eye-movement related effects.

## Introduction

The processing of faces has been a prominent object of interest in both eye-movement research examining gaze allocation on faces (Henderson et al., 2005; Hsiao et al., 2008; Yarbus, 1967; Min et al., 2015) and in EEG research examining brain activity unique to face processing (Bentin et al., 1996; Rossion et al., 2000; Rousselet et al., 2004). Whereas eye-movement research naturally allows such movements, EEG experiments were mostly done using paradigms in which eye movements were restricted to the center of the screen (e.g. Bentin et al., 1996, Rossion, 2014). In typical ERP studies stimuli are abruptly flashed at fixation point, and EEG activity is locked to stimulus onsets and averaged over trials. Such gaze-restricted paradigms reduce signal artifacts caused by the eye movements, but ignore the dynamics of natural vision and the possible role that active sampling has on perception (Martinez-Conde et al., 2004; Schroeder et al., 2010). Co-registration of eye-movements and EEG provides a powerful tool for studying visual perception in a more natural setting where participants are free to move their eyes as they wish (Dimigen et al., 2011; Nikolaev et al., 2016). Such studies examine the EEG activity locked to either fixation- or saccade-onset (Fixation- or saccade-related potentials; FRPs, SRPs) rather than to an abrupt stimulus onset.

Using SRPs or FRPs, recent studies examined neural responses in reading (Baccino & Manunta, 2005; Dimigen et al., 2011; Frey et al., 2013) visual search (Brouwer et al., 2013; Devillez et al., 2015; Kamienkowski et al., 2012; Kaunitz et al., 2014), free picture viewing (Fischer et al., 2013; Nikolaev et al., 2013),and spatial attention (Meyberg et al., 2017). Some of these studies have directly compared well known ERP effects evoked by a stimulus onset to those evoked by fixation onsets, such as the visual P1 (Kazai & Yagi, 2003), the N400 predictability effects in reading (Dimigen et al., 2011; Hutzler et al., 2007), the P3 target related response (Dandekar et al., 2012; Kamienkowski et al., 2012) or the preview effect on the N1 (Huber-Huber et al., 2019; Kornrumpf et al., 2016), generally replicating the effects using fixation related potentials. Previous FRP studies have used face images to test effects such as target detection or pre-saccadic predictions (Dimigen et al., 2009; Ehinger & Dimigen, 2018; Huber-Huber et al., 2019; Kaunitz et al., 2014), yet did not compare face-selective activity between ERPs and FRPs. A recent study (Soto et al., 2018) used a wireless mobile EEG and eye tracking system in a mock art gallery and found FRP responses differentiating face images from images of other objects. This study faces some unavoidable methodological limitations, due to its “in the wild” setup, which reduce the scope of conclusions that can be drawn. To understand categorical processing and object identification in free-viewing studies, a careful comparison of active, free-viewing FRPs, and more passive, static gaze ERPs is necessary, while controlling for low-level features, eye movement properties and activity overlap of sequential fixations, all of which were lacking, for understandable reasons, in Soto et al’s study.

Studies using ERPs consistently find that faces induce a larger negativity when compared to non-face stimuli, peaking at ∼170ms after stimulus onset with a maximum at occipito-temporal electrode sites (N170 face effect - Bentin et al., 1996; Rossion et al., 2000; Schendan & Ganis, 2013; For a comprehensive review see Rossion & Jacques, 2011). Attempts at localizing the source of the N170 face effect suggest generators in the lateral occipital-temporal cortex, in the vicinity of the posterior fusiform gyrus (Bentin et al., 1996; Di Russo et al., 2002; Itier & Taylor, 2004). Within EEG research, the N170 face effect is the prototypical evidence for category-selective response, and has been widely used to examine multiple aspects of face processing addressing cognitive, developmental and neuropsychological questions (Bentin et al., 1999; Eimer et al., 2011; Sagiv & Bentin, 2001; Towler et al., 2012; Wynn et al., 2008; Yovel, 2016). We thus concentrate here on this electrophysiological effect.

In a free-viewing scenario, the early latency of the N170 response would be especially prone to interaction with both overlap of responses from previous fixations or eye movements, as well as with robust fixation-specific visual responses. Overlapping activity from sequential fixations, which are temporally proximal, present a methodological challenge to any study of uncontrolled fixation-related activity (Dimigen et al., 2011; Nikolaev et al., 2016). Since fixation durations could be differently skewed in different conditions, FRP distortions may be systematic and appear event after trial averaging, which is commonly used in calculating ERPs. Moreover, there may be systematic differences between conditions in saccade and fixation characteristics. Some recent studies addressed this issue by including a procedure of matching eye movement characteristics such as size, direction, and duration, across experimental conditions, in order to balance out distortions (Devillez et al., 2015; Fischer et al., 2013; Kamienkowski et al., 2012; Nikolaev et al., 2016). Other studies included a training period to encourage long fixations (Kamienkowski et al., 2012; Kaunitz et al., 2014). Here, we avoided the need for fixation selection (entailing data loss and biased sampling) or participant training (entailing unnatural conditions) by adopting a multiple regression framework for de-convolving overlapping responses using the continuous data from all trials (Dandekar et al., 2012; Ehinger & Dimigen, 2018; Smith & Kutas, 2015b), similarly to techniques used in fMRI analysis.

The main goal of this study is to carefully compare face-selective activity (face vs. non-face) in active conditions, following a saccade to stimuli at the periphery, to passive conditions in which face-selective activity is evoked by an abrupt appearance of the stimuli at a fixation point, with a main focus on the N170 face effect. In both paradigms neural activity is locked to the first instance in which categorical information of the object is available^1^. However, there are important differences between passive and active sensing conditions, which could affect the measured scalp EEG components. When performing a saccade, the visual system may use both para-foveal information and information about the motor plan itself to prepare for the incoming stimulus. The temporal prediction of the self-generated saccade offset enables the system to synchronize neuronal oscillatory phase (phase reset), shown to enhance single neuron firing rates upon fixation onset even in the dark (Purpura et al., 2003; Rajkai et al., 2008). At the level of scalp EEG, it has been shown that FRPs are accompanied by a sharp positive occipital component with a peak latency of about 80 ms from fixation onset, termed the lambda response, which is much larger in amplitude than the corresponding ERP P1 peak (Kazai & Yagi, 2003). The high amplitude lambda response may be a manifestation of a phase locking neuronal mechanism, or the result of a larger change of input across the retina, compared with a more foveal change common to ERP studies, in which the stimulus is flashed at the fixation point (Thickbroom et al., 1991; Yagi, 1979). Another feature unique to active viewing is the use of para-foveal information to form expectations about the saccade target (Friston et al., 2012; Melcher & Colby, 2008). Indeed, FRP amplitudes and latencies are modulated when congruent para-foveal information is available prior to saccade initiation (Huber-Huber et al., 2019; Kornrumpf et al., 2016). When studying well-known ERP components, these fixation-related features, found both at the level of single neuron activity and in scalp EEG recordings, must be taken into consideration.

## Materials and Methods

### Participants

Thirteen healthy adults, students of the Hebrew University of Jerusalem with no reported neurological illness, participated in the experiment (Seven females and six males; age range 19-29 years, Mean=23.3 years). The participants had normal or corrected to normal visual acuity by their report. One participant was omitted from analysis due to excessive signal artifacts. Informed consent was obtained from all participants included in the study and they received either payment or class credit for their participation. The study was approved by the institutional ethics committee of the Hebrew University of Jerusalem.

### Stimuli and Experimental Apparatus

Stimuli consisted of 170 pictures of faces and 170 pictures of painted eggs, all gray-scale with equated luminance (mean pixel intensity) and contrast (Michelson contrast; (Lmax - Lmin) / (Lmax + Lmin), where L is the luminance). Faces were stripped of hair and presented in an oval aperture. Painted eggs were chosen because of their general resemblance to faces, being oval and with internal detail. All pictures were imposed on a square gray-scale clouds-like background with an additional light gray frame (RGB = [195, 195, 195]) around them (same as used in Amihai et al., 2011), which hindered identification of the objects before fixating on them. The fixation symbol at the center of the screen consisted of a light gray dot surrounded by four light gray arrows (RGB = [195, 195, 195]). Participants were seated 100 cm from the screen (Viewsonic G75f Cathode Ray Tube). Stimuli were located near the corners of the screen such that their center was 9° from the center of the fixation symbol. Stimulus size was 1.85° × 1.85° visual angles including the mask and the gray frame (Figure 1a). The screen background was a darker gray (RGB = [128 128 128]), with screen resolution set to 1024 × 768, and a 100 Hz refresh rate.

**Fig. 1.**
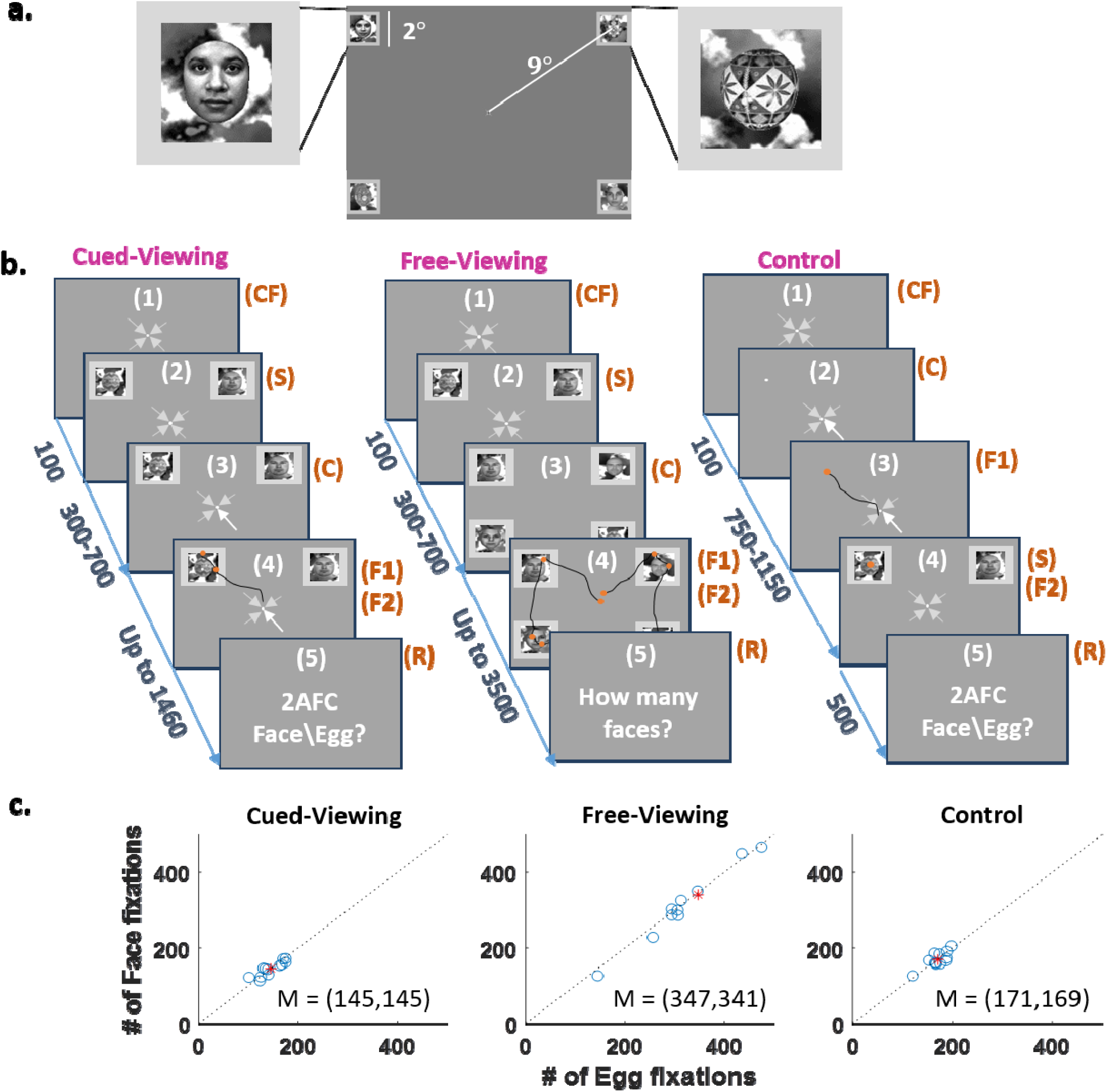
Experiment procedure. a) Two enlarged examples of a face and egg stimulus and the full screen layout preserving the correct stimulus proportions. b) Experimental design of each condition. Here the proportions of the stimuli relative to the screen are not drawn to scale to enable better understanding of the procedure. Left- The Cued-viewing condition started with a fixation on the screen center followed by stimuli appearance (S) in the periphery while center fixation is maintained (screens 1-2; the numbers in parentheses are for the sake of illustration and were not presented to the participants). After 300-700 ms an arrow cue (C) indicated the stimulus on which the participants should fixate (F1 - screen 4). After performing a saccade to the cued stimulus, participants returned their gaze to the center and reported (R) if the cued stimulus was an egg or a face (screen 5). Middle- The Free-viewing condition started like the Cued-viewing condition (screens 1-2, notice there were four stimuli in this condition as opposed to two in the Cued-viewing condition). Screen 3 depicts disappearance of the center fixation which served as a cue (C) for participants to freely view the four stimuli (screen 4). Finally, participants reported the number of detected faces in each trial (screen 5). Right-The Control condition started with a fixation on the screen center. Then, an arrow cue (C – screen 2) indicated the destination to which participants should perform a saccade (screen 3). Following the fixation (F1), stimulus appearance (S) occurred at the new fixation point (screen 4). Finally, participants reported if the stimulus was an egg or a face (screen 5). c) Mean number of first fixations on faces and eggs per participant which were included in the analysis after artifact rejection and determination of fixations of interest

### Experimental Procedure

#### The experiment included three conditions

Cued-viewing, Free-viewing, and Control (Figure 1b). All three conditions had a similar structure, but crucially, they differed in how constrained were the participants’ eye movements. In the Free-viewing condition, participants were free to explore the different stimuli at a self-driven pace and in no particular order. In the Cued-viewing condition, participants performed a controlled saccade to one of two stimuli cued by a centrally positioned arrow. In the Control condition, stimuli appeared at the fixation point. The Cued-viewing condition is an intermediate step between the completely ‘free’ viewing condition, and the passive Control condition. This provides instances of controlled saccades for which we could more carefully match the number of trials, saccade properties and control the content and timing of the fixation.

In the Cued-viewing condition, a trial started with the appearance of the fixation symbol. Once the participant maintained fixation for 100 ms (monitored by the eye-tracker), two stimuli appeared in the upper corners of the screen while participants were instructed to maintain central fixation. Between 300-700 ms after the stimulus onset, the participants received a cue informing them of the target stimulus to which they should perform an immediate saccade. The cue was the simultaneous highlighting of one of the central arrows, and a peripheral white dot in the center of the target stimulus, serving together as a combination of endogenous and exogenous cues. Stimuli remained on the screen for an additional 700-1100 ms, constraining the viewing time in each trial and promoting quick eye movement reaction to the cue. Finally, a question mark appeared in the screen center at which stage participants performed a two-alternative forced-choice task (2AFC) and indicated with a button press if the stimulus was a face or an egg.

In the Free-viewing condition, a trial started again with the appearance of the fixation symbol. Once the participant maintained fixation for 100 ms, four stimuli consisting of a random combination of eggs and faces appeared at the corners of the screen. After 300-700 ms, the center fixation symbol disappeared, cuing the participants to freely examine the four corner stimuli at their own pace and in any desired order. The behavioral task was to report the number of faces (out of the four stimuli) in each trial.

Finally, the Control condition was similar to the Cued-viewing condition, except that the endogenous (central arrow highlighting) and exogenous (white dot at the periphery) cues appeared before the appearance of the stimuli, instructing the participants to move their gaze to the white dot. After 750-1150 ms, two stimuli appeared for 500 ms at the upper corners, one of which was centered at the participant’s new fixation point on the white dot. Finally, a question mark appeared at the screen center, at which stage participants performed a 2AFC task, and indicated if the stimulus was a face or an egg. Thus, the control condition replicated the classical event-related paradigm in the sense that stimuli appeared where the eyes are fixating, but did so while controlling for a similar eye-in-head position across the Cued-viewing and Control conditions (since eyes were not pointing straight ahead as usually done with ERPs).

The three conditions were blocked, and their order was counterbalanced across participants. There were 400 trials in the Control and Cued-viewing conditions, and 300 trials in the Free-viewing condition. Trials were divided into five runs separated by a rest period. The number of faces and eggs presented in a trial were counter balanced across trials and the stimuli randomly chosen from the pool of faces or eggs, accordingly. Each condition was preceded by detailed instructions and a practice session of 20 trials.

### EEG and Eye-tracking Co-registration

EEG was acquired using an Active 2 system (Biosemi, The Netherlands) from 64 electrodes mounted on an elastic cap according to the extended 10–20 system, at a sampling rate of 512 Hz. Eight additional electrodes were placed: two on the mastoid processes, two horizontal EOG electrodes positioned at the outer canthi of the left and right eyes, two vertical EOG electrodes, one below and one above the right eye, a single EOG channel placed under the left eye, and a channel on the tip of the nose. All electrodes were referenced during recording to a common-mode signal (CMS) electrode between POz and PO3, and an online low-pass filter with a cutoff of 1/5 of the sampling rate was applied to prevent aliasing.

Binocular eye movements were recorded using a desktop mount Eyelink 1000/2K infrared video-oculography system (SR Research Ltd., Ontario, Canada), at a rate of 500 Hz. A 9-point calibration procedure, followed by a validation stage (with acceptance criteria of worst point error < 1.5°, and an average error < 1.0°) was applied before every experimental condition. A gaze drift check was performed after every 10 trials, where participants manually pressed the space bar while fixating on the screen center. The calibration procedure was repeated if an average drift of more than 1.5 degree was found. Triggers sent via a parallel port from the stimulation computer were split and recorded simultaneously by the EEG recording system and by the eye tracker, allowing offline synchronization between the records. Eye position and pupil data generated by the Eyelink system were also recorded by the Biosemi system as additional analog channels. This was used to ensure correct synchronization but the analysis was performed on the digital data recorded by the eye tracker, which has a better signal-to-noise ratio (SNR).

### Eye Movement Data Analysis

Saccades and fixations were detected using the Eyelink proprietary algorithm, which identifies when the moment-to-moment velocity and acceleration of the eye exceeds a pre-determined threshold velocity of 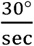, an acceleration of 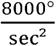, and a minimum position change threshold of 0.1°.^2^ Analysis was performed on the participants’ dominant eye. Exclusion criteria for detected saccades were: 1) If a blink occurred during the saccade, 2) If the saccade amplitude exceeded the diagonal screen size (> 28°) or 3) If the saccade duration was longer than 100ms, which is unlikely. Fixations were excluded if their duration was longer than 2.5 sec, if they occurred within 20 ms of a blink, or if they followed an invalid saccade according to the above criteria. We verified that the difference between a detected saccade offset and its ensuing fixation onset was no more than 4 ms. On average, ∼1.2% of fixations were removed per participant due to these exclusion criteria. These events were kept for consideration in the regression analysis (see Fixation Related Potentials section), but were marked as invalid events. Blinks were defined from 5 ms before the eye tracker lost gaze location to 5 ms after gaze was reestablished.

Detection of fixation onsets using either the Eyelink algorithm, or the statistics of eye-movement velocity using the algorithm published by Engbert & Kliegl, 2003, are based on some pre-defined parameters which might not be optimal for every fixation in the data. As opposed to saccade onsets, fixation onsets are especially challenging since saccades terminate with different motion characteristics such as undershooting (or corrective saccades; Prablanc & Jeannerod, 1975; Kapoula & Robinson, 1986) and post-saccadic oscillations (Hooge et al., 2015; Tabernero & Artal, 2014). It is not clear what is the optimal algorithm for fixation onset detection under different experimental conditions (Andersson et al., 2017). We performed an attempt to realign the fixations based on the local movement statistics around each fixation separately. A window of ±100 ms around the fixation onset, determined by the Eyelink algorithm, was defined and further constrained to include only one preceding saccade. We then corrected the fixation onset to be the first point to cross below two standard deviations of the eye velocity data. This resulted in realigning the fixations to the end of the main saccadic movement (see Supplementary Figure 1). The FRPs were hardly affected by the different onset definitions (data not shown). The same procedure was also done for fixations defined by the Engbert & Kliegl, 2003, algorithm, with similar results.

Fixations of interest (FOIs) were detected within the stimuli region of interest (ROI). The ROI was defined as the area within the external frame of the object picture (Figure 1). We verified that fixations within this frame were sufficient for the participants to accurately report the object identity (Supplementary Figure 2). A valid “first fixation” (F1) in the Cued and Free-viewing conditions was the first fixation within the ROI of a stimulus which followed a saccade larger than at least ∼6° visual degrees. This ensured that a previous fixation, if happened, was far enough from the stimulus to hinder para-foveal identification. Fixations occurring within 20 ms from a blink, or during EEG artifacts, were marked as invalid fixations and were modeled separately. Overall 7% of fixations were considered invalid.

The final number of fixations on faces and eggs included in the analysis after rejection for any reason (including EEG artifacts) remained balanced (Figure 1c). The Free-viewing condition naturally resulted in an overall larger number of fixations per participant and more variability between participants due to the less controlled task and the two additional stimuli presented per trial. The total number of observations used in the Cued-viewing condition was 1742 fixations on eggs (mean=145 per participant), and 1735 fixations on faces (mean=145 per participant). The Free-viewing condition included a total of 4167 fixations on eggs (mean=374 per participant), and 4096 fixations on faces (mean=371 per participant). The Control condition included a total of 2048 fixations on eggs (mean=171 per participant), and 2033 fixations on faces (mean=169 per participant). In the Cued-viewing condition saccade amplitudes of the fixations used in the analysis (first fixations on stimulus) were distributed normally with a mean and standard deviation of *M* = 8.33°, *σ* = 0.83, consistent with the distance of the stimuli from the center fixation. In the Free-viewing condition saccade amplitudes were fitted to a mixture model of two Gaussians with *M*_1_ = 8.96, *σ*_1_ = 0.81, *M*_2_ = 14.23, *σ*_2_ = 0.65, due to the different distance between horizontally and vertically adjacent stimuli. Fixation durations were skewed right with a median value of 202 ms and 218 ms in the Cued and Free-viewing conditions respectively and a maximum of 1600 ms (see Supplementary Figure 3).

### EEG Pre-processing

Data processing was done with custom MATLAB code (Mathworks, Natick, MA), with some functionality adopted from the Fieldtrip toolbox (Oostenveld et al., 2011) and the EEGLab toolbox (Delorme & Makeig, 2004). Electrodes that included excessive noise across the entire experiment duration (based on visual inspection) were excluded (1-2 channels in four out of 12 participants). Next, data was re-referenced to an average of all electrodes excluding the eight most frontal electrodes (Fp1,2,z and AF7,3,z,4,8), which are sensitive to eye movements. Since we pause the recording during breaks between blocks, amplitude steps at the onsets of recording could be smeared later by high pass filtering. To avoid this, we performed linear de-trending within each block by subtracting the linear vector connecting the means of the initial and final 10 samples of the block, which enabled us to treat the complete dataset as one continuous recording. The data was high-pass filtered with a cutoff of 0.1 Hz with a third-degree, zero phase-shift Butterworth filter, and 50 Hz line noise was removed with a custom notch filter designed to suppress continuous 50 Hz oscillations, typical of power-line noise, while having less effect on more transient components (Keren et al., 2010). To attenuate noise driven by eye movements and muscle activity, independent component analysis (ICA) was applied. ICA training data was generated by concatenating segments of −500 to +1000 ms around stimuli onsets, as well as −30 to +30 ms segments centered on every saccade-onset event (Keren et al., 2010). ICA components were manually identified as reflecting ocular noise based on their temporal profile, scalp topography, power spectrum, and low-frequency power modulation around blinks, saccades or stimulus onsets (on average, 11 components were selected per participant, range = [7, 13]). To remove noise components, these components were multiplied by zero and the data was then re-mixed into channel data. Next, segments exceeding a threshold of ±100 µV were marked as artifact and excluded from analysis together with a margin of 40 ms around the artifacts, following verification with visual inspection. Finally, electrodes excluded before pre-processing were recreated by mean interpolation of the neighboring electrodes.

### Fixation-Related Potentials

As mentioned in the introduction, a challenge of analyzing EEG data aligned to fixation onsets is both temporal overlap of brain responses evoked by consecutive fixations and the effects that saccade properties have on this post saccadic neural response (Dimigen & Ehinger, 2019). To address these challenges, a multiple regression model was applied directly to the continuous EEG data, in order to disentangle (de-convolve) the overlapping event-related responses to multiple events occurring in a semi-controlled manner in each trial (Smith & Kutas, 2015a, 2015b). We defined four temporally-adjacent events within a trial, for which we wanted to model a separate waveform: 1. Stimulus appearance (S); 2. Cue appearance (in the Cued-viewing and Control conditions) or disappearance (in the Free-viewing condition) (C); 3. First fixation on the stimulus (or the white dot in the control condition) (F1); 4. Second immediate fixation on the stimulus (F2). The sequence of events per trial for each condition (see Figure 1) was therefore: *S* → *C* → *F*1_*E\F*_ → *F*2_*E\F*_ for the Cued-viewing and *S* → *C* → *F*1_*E\F*_ → *F*2_*E\F*_ … → *F*1_*E\F*_ for Free-viewing condition, and *C* → *F*1 → *S*_*E\F*_ → *F*2_*E\F*_ in the Control condition (*E\F* stands for eggs and faces – these were considered separate events in the model). An additional event accounted for in the model was a “general fixation”, that is, any fixation which did not meet the criteria of FOIs (see Eye Movement Data Analysis section). The response evoked by each event was modeled time-point by time-point over a window of 2000 ms (−200 to +1800 ms^3^) sampled at 512 Hz, which resulted in a total of 1024 data points. We defined dummy coded predictors (or stick functions) *X*_*event n, latency l*_ that are 1 for data points measured at a specific latency *l* relative to the time-locking event *n* and 0 otherwise. In our model we therefore have *l* = 1024 data points and n = 7 events of interest resulting in a total of 1024*7 predictors in the model. If *l*_2_ is an index that runs on all latencies of interest around an event onset, the predictor for a specific latency *l*_1_ will be 1 only when *l*_1_ = *l*_2_ and zero otherwise (Equation 1).

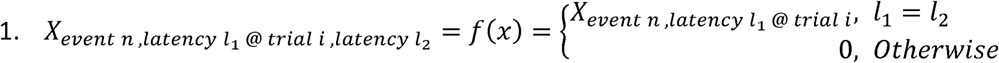

Thus, each measured data point is modeled as a sum of the effects of overlapping event-related time series (Equation 2):

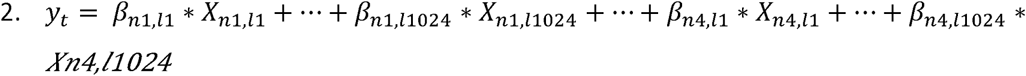

For example, for a series of events in the Cued-viewing conditions (Figure 1), voltage *y* recorded 600 ms past stimulus onset (S), 300 ms past arrow cue onset (C), 100 ms past first fixation onset (F1), and before a second fixation (F2) the regression equation would be (Equation 3):

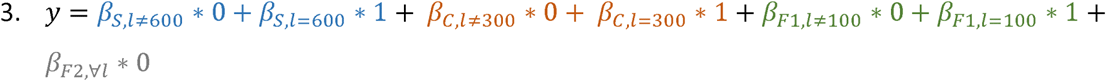

The analysis was carried out in MATLAB (except for the regression stage which was run in Python, which handled the memory load required more efficiently than MATLAB^4^).

In addition to the binary event regressors, we included in the model a continuous-variable predictor of the saccade amplitude preceding a fixation, expanded over five spline basis functions (Ehinger & Dimigen, 2018), to account for the non-linear effect that saccade size has on the lambda peak (Dandekar et al., 2012; Dimigen et al., 2009; Ries et al., 2018). We removed from the continuous data (thus from the design matrix) any data points defined as artifacts. The regression procedure resulted in ‘regression ERPs\FRPs’ (rERPs in the Control condition, rFRPs in the Cued and Free-viewing conditions), which are the time-series of estimated coefficients per event. rERPs\rFRPs were then averaged over participants and baseline-corrected by subtracting the average of 200 ms before the event of interest. For convenience going forward, we drop the ‘r’ prefix and refer to FRPs and ERPs. (For an additional explanation of the multiple linear regression model and the spline basis method for modeling saccade amplitude, see Dimigen & Ehinger, 2019).

In the Free-viewing condition participants freely viewed different combinations of face and egg stimuli, and hence the fixation on a stimulus could be affected by which stimulus was previously fixated. We thus created a second model, only for the Free-viewing condition, in which the F1 event was divided into four sub-events according to the combination of pre and post fixation categories. There were four such combinations: Face->Egg, Egg->Egg, Egg->Face, and Face->Face. Thus, the full model included n = 9 events of interest: *S* → *C* → *F*1_*EF\FE\EE\FF*_ → *F*2_*E\F*_ … → *F*_*general*_. A total of 2023 fixations within categories (Egg->Egg or Face->Face, per participant: mean=170, range=[71, 258]) and 989 examples of fixations between categories (Face->Egg, or Egg->Face, per participant: mean=83.5, range=[25 120]) (Figure 3), were used in this regression analysis.

**Fig. 2.**
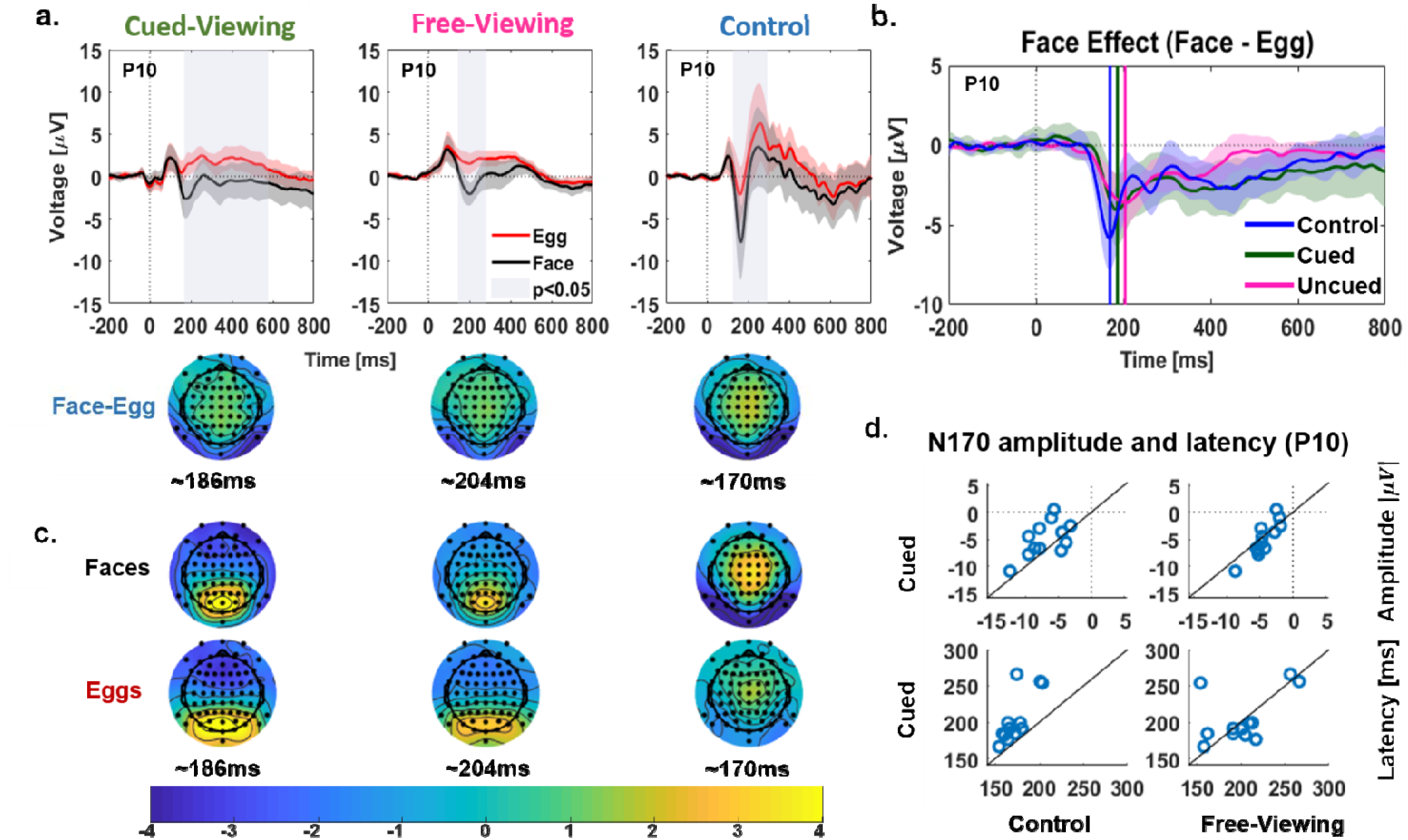
Comparison of the ERP and FRP face effect. a) Top: the grand average regression ERPs/FRPs obtained by egg and face stimuli separately for each experimental condition shown for electrode P10. Waveforms include a 95% confidence interval. Windows of significant difference marked based on a time and space cluster permutation statistics for p<0.05. Bottom: Scalp distributions of the face effect averaged over a window of 30 ms around the N1 peak latency per condition. b) Faces-eggs difference wave in all three conditions superimposed with a 95% confidence interval (shaded error-margins) and the peak latencies (vertical lines). c) Scalp distributions of the activity elicited by faces and eggs separately, averaged over a window of 30 ms around the N1 peak latency per condition. d) Single subject N170 face effect peak amplitudes (top) and latencies (bottom) compared between conditions – Cued-viewing vs. Control (left) and Cued vs. Free-viewing (right)

**Fig. 3.**
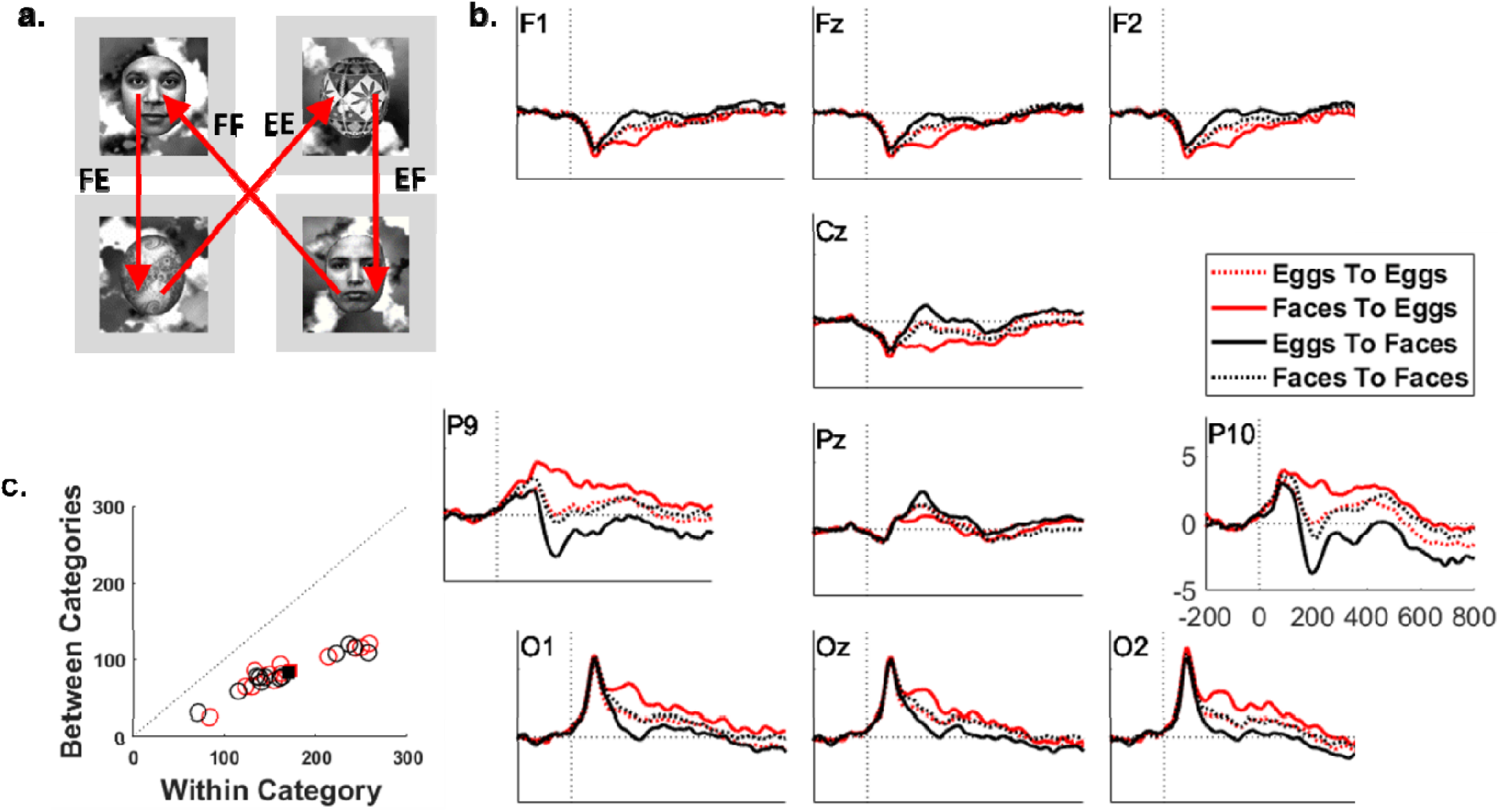
a) Illustration of the four different categories of fixations, depending on the previous fixation (FF – face following face, EF – face following egg, EE – egg following egg, FE – egg following a face. b) Grand average FRPs per combination of pre-post fixation categories for a subset of the electrodes. c) Number of fixations of each stimulus type (red-egg, black-face) for within and between category fixations. Due to the configurations of stimuli there were more cases of within category fixations than between categories.

### Statistical Analyses

We first examined the a-priori defined N170 face effect. To that end, we determined the peak of the face-egg difference activity in electrode P10, for each participant. The peak was defined by finding the local minima within a time window of 120-280 ms (large window around the N1). A one-way repeated measures analysis of variance (rmANOVA) on the amplitudes and latencies of the N170 face effect peaks was conducted across experimental conditions, followed by planned comparisons between ERPs and FRPs (Control ERP vs. the average of the FRP conditions), and between the two FRP conditions.

To test for significant differences in activity elicited by faces vs. eggs beyond the a-priori defined N170 time-window, we used a cluster permutation algorithm (Maris & Oostenveld, 2007) during which a two-tailed t-test was conducted for each time point and electrode separately for the three conditions. t-values that crossed a threshold of p < 0.05 were chosen and clustered on the basis of temporal and spatial adjacency. For each cluster, the sum of t-values (t_sum) was determined. To define the distribution of the t_sum under the null hypothesis of no differences, this procedure was iterated, where in each iteration the assignment of data to face or egg was permuted covering all 2^12^ possible permutations, and the maximal t_sum value was recorded in each iteration. Finally clusters in the original analysis with t_sum > the 95% percentile of t_sums in the null distribution were considered as significant.

To test for statistical differences of the face effect between ERPs and FRPs, we ran a time-resolved one-way rmANOVA on the face-egg difference activity across the three experimental conditions (in effect examining the interaction between stimulus type and experimental condition). We tested this using the cluster permutation method over the F-statistic (2^12^ permutations, p <0.05). The ANOVA was followed with two planned comparisons: between the ERP condition (Control) and the FRP conditions (collapsed over the Cued and Free-viewing), and between the two FRP conditions. Both of these comparisons were again done using the cluster permutation t-test algorithm.

### Post-hoc Test of Para-foveal Categorization

To verify that para-foveal categorical information is indeed limited when fixating centrally using the screen layout and stimuli, a behavioral experiment was conducted (including 12 participants of mean age = 25.7) during which participants either determined the category of a cued stimulus or counted the number of faces presented while they maintained a center fixation within a 1.5° radius (verified by eye tracking). Presentation parameters were similar to the ones used in the main experiment. A binomial test indicated that 10 out of 12 participants performed at chance level (p>0.05, 1-sided) in the Free-viewing condition and that 9 of 12 performed at chance level in the Cued-viewing condition. The few participants who showed significant behavior performed only slightly above chance and were not the same participants in the two conditions, suggesting that, even in these participants, the above chance performance does not indicate robust para-foveal recognition (Supplementary Figure 2).

## Results

### Behavioral Results

Participants were highly accurate in performing the behavioral tasks: detecting faces or eggs in the Control and the Cued-viewing task (control: hit rate M = .88, SD = .05; Cued-viewing: M = .91, SD = .03), and counting the number of faces in the Free-viewing condition (correct count M = .85, SD = .09). We ran a rmANOVA on the accuracy scores to test for differences between the conditions, and followed it with post hoc comparisons. The result in the ANOVA was marginally significant F(2,22)=3.3827, p=0.037. All three t-test comparisons between the conditions were not significant after Bonferroni correction for multiple comparisons.

### FRPs Compared to ERPs

The main goal of this study was to compare the free-viewing N170 face effect to the classical N170 face effect in time and space. We thus compared activity for faces vs. eggs (our non-face control stimulus) in three different conditions: Cued-viewing, Free-viewing, and Control. Note that in all three conditions, the gaze direction was deviated relative to the head. We first verified that we obtained the N170 face effect in our Control condition. Indeed, we observed a significantly larger negativity for faces compared to eggs peaking around 170 ms. The topography of the N170 face effect (the difference between faces and egg responses) was also consistent with previous studies, with a maximum at lateral temporo-occipital electrodes (Bentin et al., 1996) and a concomitant frontal positivity (vertex positive potential or VPP (Joyce & Rossion, 2005; Jeffreys & Tukmachi, 1992; Figure 2). Critically, in both of the FRP conditions (Cued and Free-viewing), we obtained a clear N170 face-specific effect, namely, the response to faces was significantly more negative than eggs at the expected latency of around 170 ms and with a typical topography including positivity in the fronto-central electrodes, and negativity in the temporal-occipital electrodes. (Figure 2a, Supplementary Figure 4). In all three conditions, the significant spatio-temporal clusters extended for additional 200-300 ms beyond the N170 peak.

### Planned Comparison of the N170 Face Effect Across Conditions

The peak of the N170 face effect was smaller in amplitude and later in time for the FRPs (Cued and Free-viewing conditions) relative to the ERPs (Control condition – Figure 2b). A one-way rmANOVA of the N170 face effect peak amplitude across conditions showed a significant effect of condition (F(2,22) = 9.474, p = 0.001). We followed it up by planned comparisons of the ERPs (Control condition) against the FRPs (Cued and Free-viewing conditions): the ERP N170 face effect in the Control condition elicited a larger negativity than the two FRPs combined (t(11) = 3.66, p = 0.003), while the two FRPs did not significantly differ (t(11) = 1.37, p = 0.2). Figure 2d depicts these differences by comparing the individuals’ peak amplitudes in the Control vs. Cued-viewing conditions (top left) and the Cued vs. Free-viewing conditions. While most participants show larger peak amplitudes in the Control condition than the Cued-viewing condition, the amplitudes between the Cued and Free-viewing are mostly similar (top right). A similar rmANOVA for peak latency of the N170 face effect found a significant effect of condition (F(2,22) = 7.32, p = 0.004); follow up tests showed that the latency of the ERP was significantly earlier than that of the two FRPs combined (t(11) = −4.3, p =0.001), while the difference between the two FRP latencies was not statistically significant (t(11) = 0.2, p = 0.84). Again these differences can be seen in participants’ individual data (figure 2d-bottom). The differences in amplitude and latency remained similar whether we used the fixation times based on the propriety Eyelink definitions or if we further realigned fixation onsets (see Eye Movement Data Analysis section).

### Exploratory Analysis

To explore face effect (face-egg) in wider spatio-temporal window, we conducted a one-way ANOVA on the difference wave, in all electrodes and time points in the range of [0 800] (thus looking at the interaction between stimulus type and experimental condition). The cluster permutation test, using a dependent sample F-statistc, revealed one marginally significant cluster (p=0.03) in a central region emerging at a latency of ∼400-550 ms. We followed this up with a planned comparison between the ERP (Control condition) and FRPs (the average of the Cued and Free-viewing conditions), and between the two FRP conditions. When the analysis was done across the entire 800 ms epoch, no significant clusters were found. When the analysis was restricted to the time-window of the significant cluster from the ANOVA, there was a significant cluster indicating a more positive response in the Cued-viewing vs. the Free-viewing conditions (Supplementary Figure 6). Thus, although there were some differences in the later time windows, they require further confirmation.

### Adaptation Effects

Since the task in the Free-viewing condition was to freely explore four stimuli sequentially, as opposed to one per trial in the Cued-viewing and Control conditions, we could explore the N170 adaptation effect (Amihai et al., 2011; Cao et al., 2015; Feuerriegel et al., 2015; Kloth et al., 2010; Kovács et al., 2006; Maurer et al., 2008), by comparing FRPs of first fixations on stimuli following saccades from a stimulus of the same vs. different category (Figure 3): Face->Egg, Egg->Egg, Egg->Face, and Face->Face. Notably, the responses to faces and eggs shifting within category (i.e. face following face, and egg following egg) were midway between the responses to faces and eggs for between-category fixations (i.e. face following egg and egg following face). That is, fixations on faces after viewing an egg elicited the most negativity ∼200 ms after fixation onset while fixations on faces following other faces elicited an attenuated negativity, as expected in an adaptation framework. Similarly, fixations on eggs following faces elicited the largest positivity ∼200 ms in occipito-parietal electrodes and negativity in frontal electrodes, whereas when following other eggs, the response amplitude was attenuated. As a consequence, the most extreme difference between the response to faces and eggs (the face effect) was obtained when the gaze shifted between categories, whereas the face effect was strongly attenuated when the gaze shifted within category.

### Scalp Distributions

The scalp topography of the N170 face effect at its peak latency showed similar distribution across conditions (Figure 2a,b). However, this was not the case for the responses to faces and eggs separately (Figure 2c). Broadly, for both faces and eggs, fixation-related responses were dominated by a midline occipital positivity and concomitant frontal negativity, whereas event-related responses showed a central-frontal positivity accompanied by occipito-temporal negativity. Next, we examine these differences in more detail.

Figures 4 and 5 depict the evolution in time of the activity elicited by eggs, faces and the face-egg difference, for each condition. In general, the topographies of the responses to each category in the Control condition are similar to the topography of the differential face-egg response, with a clear N1 peak (∼170 ms) in the lateral temporo-occipital electrodes, and a positive counterpart in the fronto-central electrodes (vertex positive potential, VPP). The first difference between the Control and both of the active viewing conditions, when examining within category responses, emerges during the P1 peak (100 ms) (Figures 4, 5). While the positive voltage at lateral occipito-temporal electrodes (P9\P10) is evident for all conditions (Figure 2a, Figure 4), a central occipital positivity starting as early as 80 ms and accompanied by a frontal negativity is seen for the FRPs, but not for the ERPs. This occipital positivity is maintained also during the N1 time-window such that the FRPs show no N1 in medial occipital electrodes nor its positive counterpart in frontal electrodes. However, occipital-temporal (P9\P10) electrodes show a clear N1 deflection (∼190 ms) in these conditions. Taken together, these results show that while the category selective activity (face-egg) is topographically similar whether locking the neural signal to stimulus onset or fixation onset, the overall neuronal signals elicited by each category on its own are different between the two viewing conditions.

**Fig.4.**
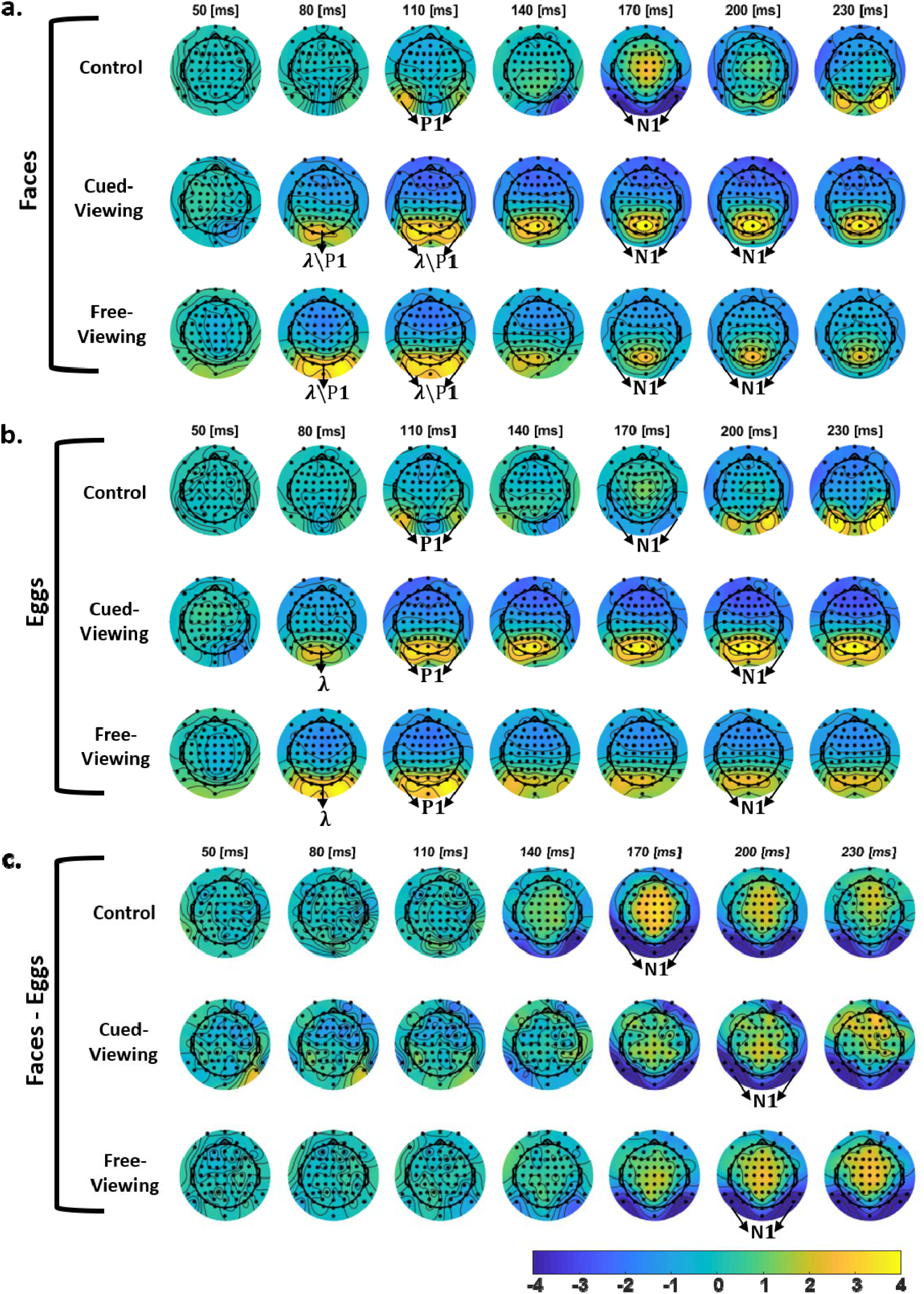
Development in time of the response to faces (a), eggs (b) and to the face-egg difference wave (c). The Lambda response in the FRPs starts earlier than P1 of the ERP and is more “spread out” including central occipital electrodes. The occipital component is sustained in the FRPs. N1 peaks later in the FRPs

**Fig.5.**
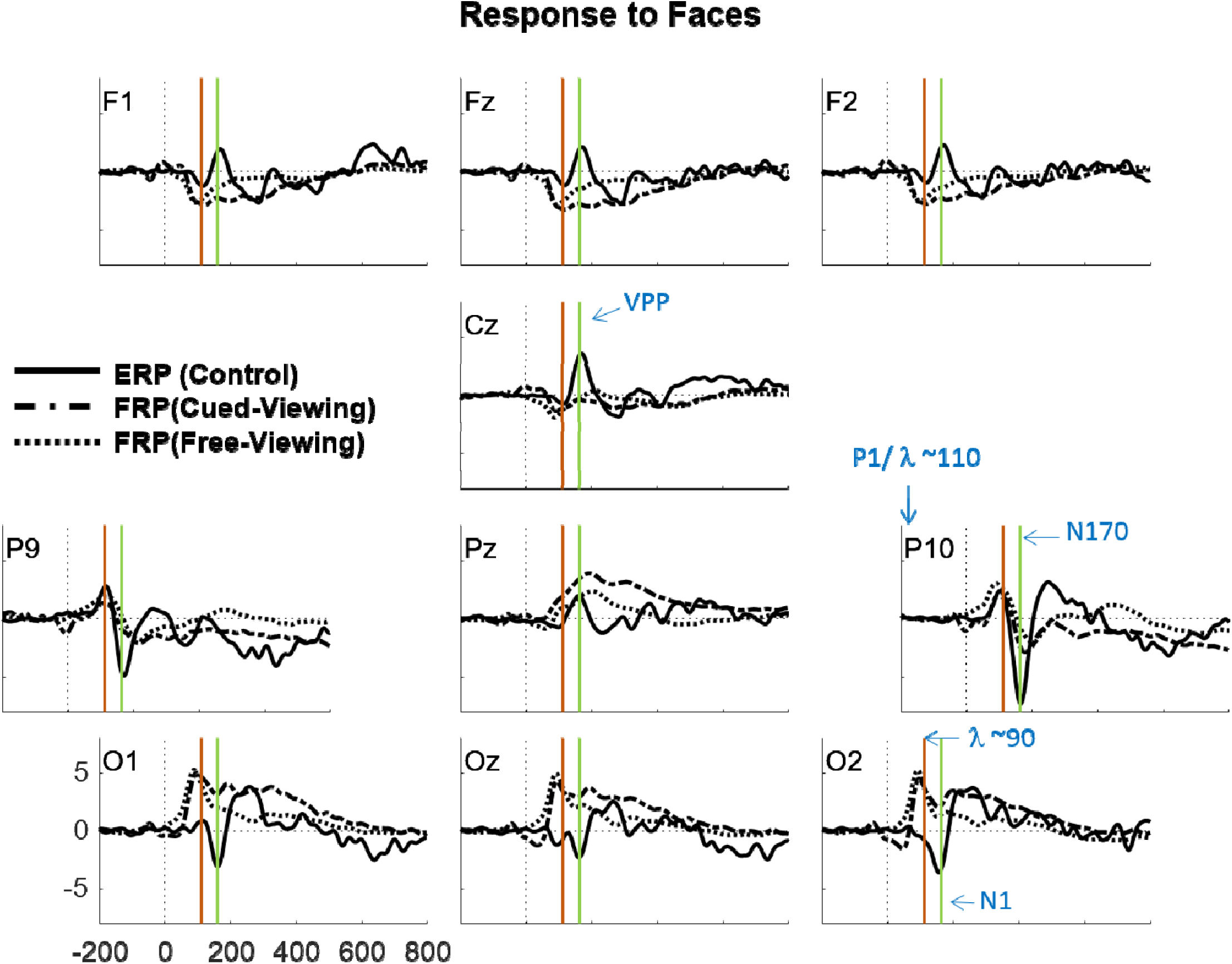
ERPs\FRPs of the response to faces in a subset of the electrodes. The red and green vertical lines indicate the P1 and N1 peaks respectively. While the Central and Parietal electrodes (P9\10, Cz, Pz) show similar waveforms, the main differences emerge in the Occipital and Frontal electrodes (F1\z\2, O1\z\2). For similar results in response to eggs, see Supplementary Figure 5.

What could be the reason for the striking difference in the scalp recorded responses between ERPs and FRPs, a difference that nevertheless preserves a similar face effect? One explanation could be that the initial P1\lambda response, which is different for a stimulus versus a fixation onset (Yagi, 1979), may overlap in time with the categorical processing reflected by the occipito-parietal N1. To test the plausibility of this hypothesis, we simulated the development in time of possible dipole sources using the Brain Electrical Source Analysis (BESA) software (MEGIS Software GmbH, Gräfelfing, Germany). Note that this simulation was not an exploratory source localization (“inverse solution”) analysis, but rather a “forward” solution designed to test the plausibility of a hypothesis. That is, we seeded dipoles in hypothetical locations and adjusted them within the constraints of previous dipole localization literature, and with hypothetical time courses, and examined the resulting scalp distribution over time. First, the face effect was modeled by two transient dipoles starting around 90 ms and peaking around 170 ms, located in right\left occipital-temporal cortex (roughly Brodmann areas 37 of the posterior fusiform gyrus; Bentin et al., 1996; Di Russo et al., 2002; Itier & Taylor, 2004; Soto et al., 2018). Next, we added two lateralized dipoles modeling the P1 activity located in the visual association area in the extra-striate cortex (roughly Brodmann 18\19; Clark et al., 1995; Mangun, 1995; Martínez et al., 1999) oriented towards the midline, starting around 60 ms and peaking around 100 ms with slowly decaying sustained activity (Figure 6a-b). This seems plausible as Di Russo et al., 2002, have reported an overlap of the N1 and P1 sources in visual evoked potentials. The simulation configuration of four dipoles peaking at two different time points simulated the ERP topography in the Control condition (Figure 7). The difference between the ERP topography and the FRP topographies could be captured by adding a single dipole over the Calcarine sulcus with posterior\anterior orientation (Figure 6), previously suggested as the location of the lambda response generator, which is activated upon a new fixation (Kazai & Yagi, 2003). This additional dipole causes a reversal in the FRP components in the occipital and frontal electrodes compared to the control ERPs, hardly affecting the FRP component orientations in occipital-parietal electrodes (P9/P10) (Figure 7), as seen in the experimental data.

**Fig.6.**
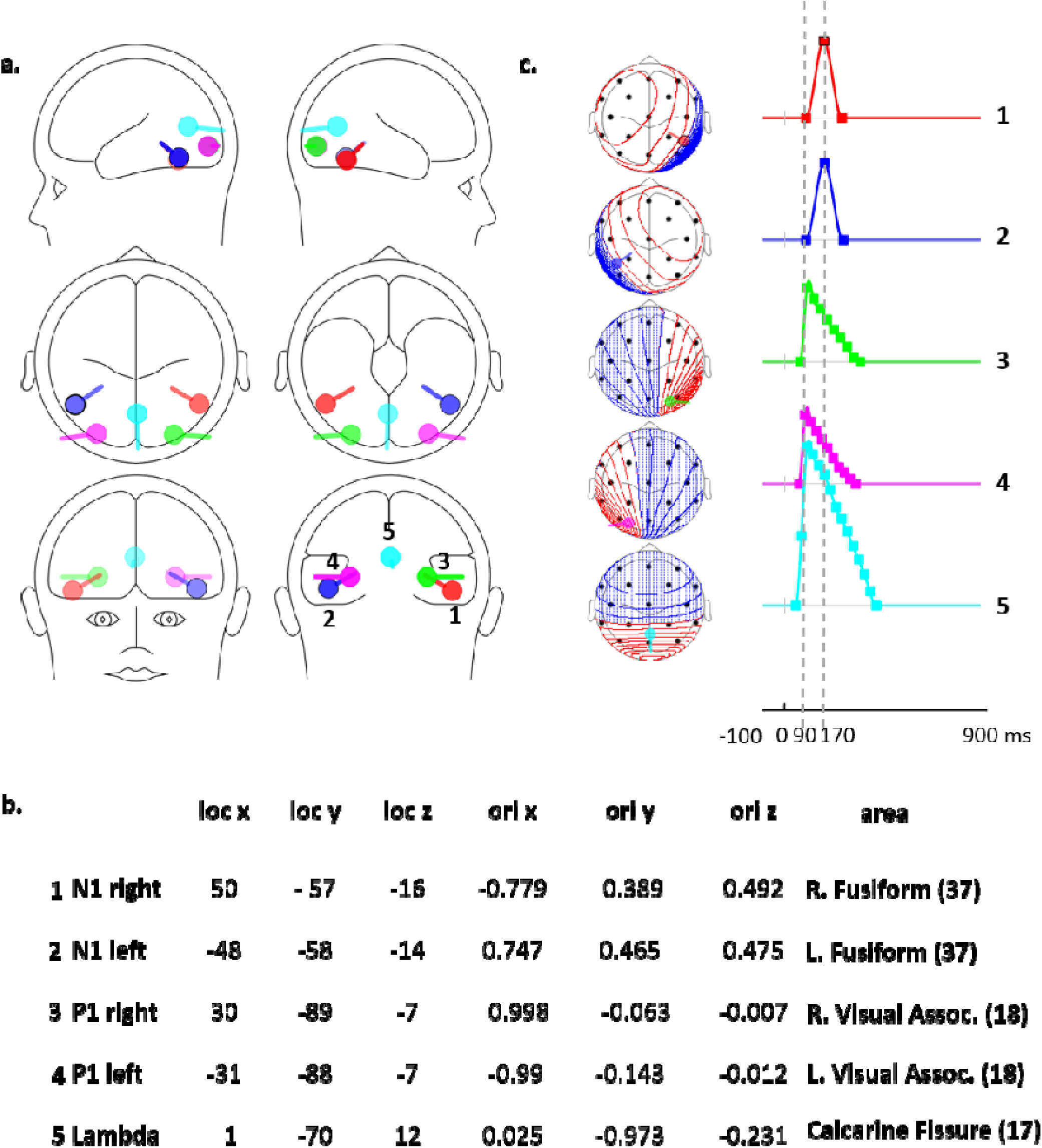
Source model using BESA simulator. a) The Spatial organization of the dipoles used in the model. b) Position of the modeled dipoles in Talairach coordinates. c) The dynamics of each dipole. P1\lambda dipoles have a nonsymmetrical dynamics and peak at 90 ms, N1 dipoles peak at 170 ms.

**Fig.7.**
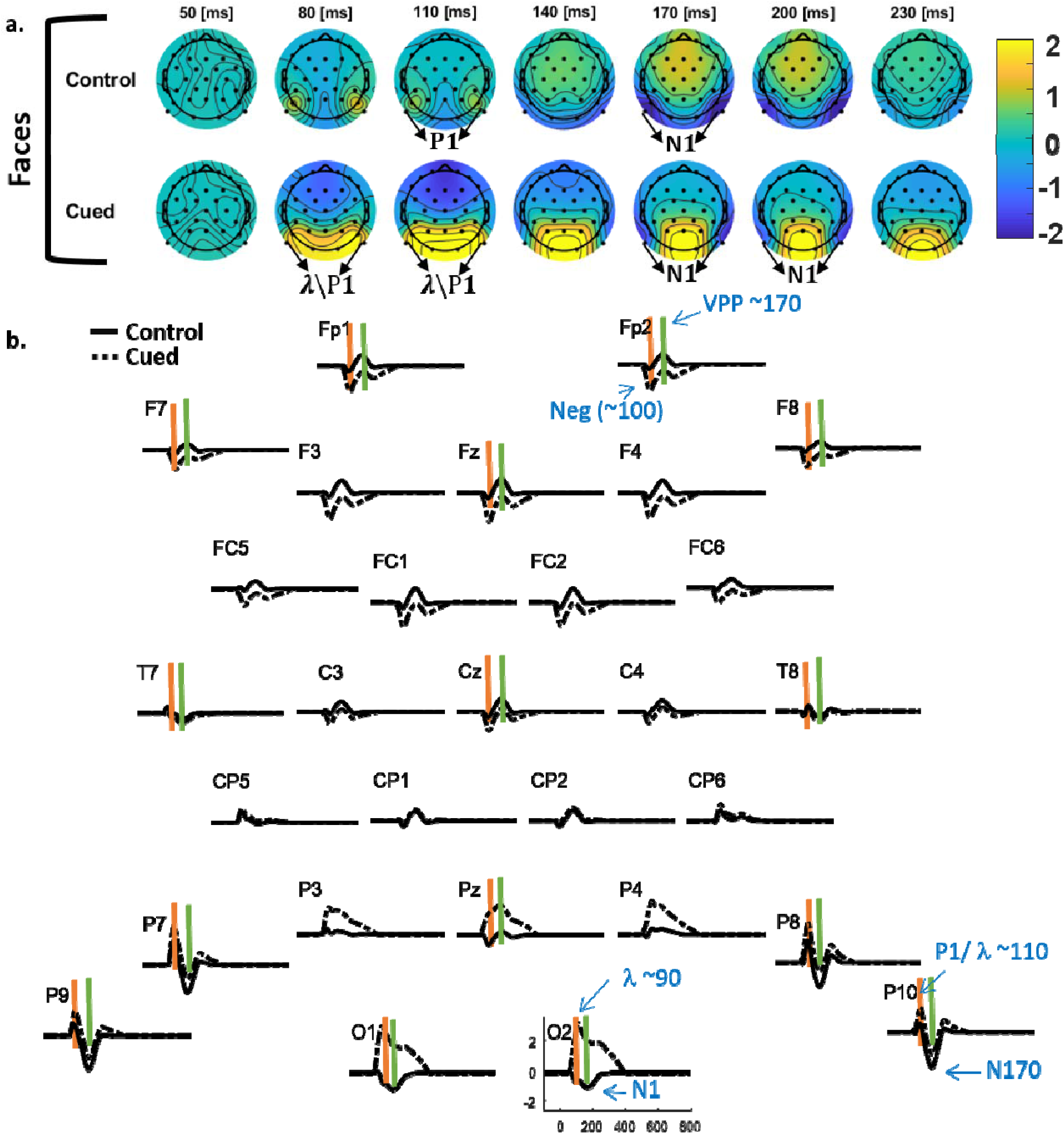
Simulated topographies (a) and waveforms (b). The classic control ERP is modeled using four dipoles (Figure 6) reflecting the P1 and N1. The Cued-viewing condition FRP is modeled by adding the “lambda” dipole. The red and green vertical lines indicate the P1 and N1 peaks respectively as in Figure 5.

## Discussion

Traditional studies of visual perception rely mainly on a serial presentation of stimuli at the center of the visual field, while the viewer tries to keep his or her gaze fixed. Such well-controlled paradigms have obvious methodological benefits, but they are quite removed from real-life perception. In real-life, stimuli do not appear abruptly out of the void, and viewers are free to search and scan in a self-paced manner, as well as benefit from background (peripheral) information. The overall goal of using fixation-related, rather than event-related, potentials is to study visual cognition while taking into account the dynamics of natural vision and the possible role that active sampling has on perception (Dimigen et al., 2011; Nikolaev et al., 2016). The focus of the current study was on a well-known signature of category selective processing, the N170 face effect, and its manifestation in fixation-related potentials, compared to the traditional way of presenting stimuli serially at fixation. This is an essential step towards bridging the gap between perception in more natural viewing, and what we know about categorical perception from artificial laboratory experiments. We demonstrated that the N170 face effect, that is, the difference between the response to faces and non-face stimuli at a latency of ∼170ms, is clearly apparent when locking EEG activity to fixation on a peripheral stimulus, with a similar topography to that found using ERPs generated by stimulus onset at fixation point. This validates the use of the N170 face effect as a signature of face (vs. non-faces) processing in free-viewing experiments.

### N170 Face Effect in ERPs and FRPs

The N170 face effect peak had somewhat of a smaller amplitude and longer latency later in the FRP active viewing conditions compared to the ERP passive viewing condition. Smaller FRP than ERP amplitudes are consistent with evidence from neuronal activity in the visual cortex of non-human primates’ (areas V1, V2, V4, and LIP), showing a reduced firing rate modulation locked to fixation onset, when compared to that evoked by a flashed stimulus (Gallant et al., 1998; Kusunoki et al., 2000, but see: MacEvoy et al., 2008). A somewhat late N170 face effect (peaking at 186 ms, like in our Cued-viewing condition) was also found by Soto et al., 2018, in a free-viewing situation, although this study had no comparison to ERPs.

The differences in amplitude and latency may have both mechanistic and methodological sources. First, in the ERP case, the onset of the stimulus is abrupt, whereas the onset of fixation is less discrete and could entail a gradual onset. This could result either from gradual release from saccadic suppression, or the incremental accumulation of information as the eye slows down to a halt. An abrupt change of input in neuronal receptive fields is likely to create stronger and more immediate activation of neurons and could lead to larger measured responses. In addition, beyond their immediate effect on visual processing, abrupt stimulus onsets are known to capture attention in an exogenous manner (Carrasco, 2011; Egeth & Yantis, 1997; Yantis & Jonides, 1984) and subsequently increase the gain of neural activity (Busse et al., 2008). Second, an important difference between passive and active viewing conditions is the utilization of para-foveal information in active viewing paradigms, which is unavailable when a stimulus is flashed from the void. Para-foveal processing is an essential part of visual processing in natural conditions (Dimigen et al., 2012; Henderson & Anes, 1994; Rosenholtz et al., 2018; Schotter et al., 2012), providing cues for predicting the target of the next fixation. Co-registration studies directly addressing the question of the para-foveal processing effects using faces revealed earlier N1 latencies and smaller amplitudes for valid vs invalid para-foveal information (Buonocore et al., 2019; Huber-Huber et al., 2019). In our experiment, para-foveal information was extremly limited due to the small size of the sitmuli, their peripheral location, and the presence of a surrounding mask. A behavioral test of covert stimuli categorization revealed that participants performed at or around chance level (Supplementary Figure 2). Since para-foveal expectations should shorten the FRP N1 latencies, this cannot be the reason for our longer FRP latencies compared to ERPs. However, we cannot rule out the possibility that rudimentary para-foveal information could lead to variable expectations across trials.

From a methodological point of view, the inherently less-determined moment of fixation onsets, compared to the well-determined timing of the stimulus onset, introduces potential trial-by-trial variability in FRPs. This can smear the estimated waveform and lead to the smaller amplitudes and later latencies of the FRPs compared to ERPs. There are numerous algorithms for the detection of different types of eye-movements, including fixations. Andersson et al., 2017, compared ten such algorithms to “human-expert” definitions of saccades, fixations, smooth-pursuit, and post-saccadic oscillations. The study reveals quite dramatic variability between the eye movement characteristics defined by the different algorithms, presenting an important warning sign. Different algorithms seem to perform more “human like” for different types of eye movements. However, since there are no theoretically motivated thresholds for determining the oculomotor parameters of each eye-movement type, it is not even clear that the human-experts’ data labeling is a reliable gold standard. Fixation onsets, as opposed to saccade onsets, are especially challenging since saccades terminate with different motion characterisics such as immediate corrective saccades (Kapoula & Robinson, 1986; Prablanc & Jeannerod, 1975) and post-saccadic oscillations (Hooge et al., 2015; Tabernero & Artal, 2014). In this study we analyzed the data once with fixation onsets defind by the Eyelink proprietary algorithm (which is based on predetermined thresholds on the velocity, acceleration, and position change), and again after realigning the fixations to the end of the saccade using very local statistics around each saccade separately (see Eye Movement Data Analysis section and Supplementary Figure 1). These attempts had almost no effect on the results, suggesting that either we did not manage to reduce the inter-trial variability in this way, or that such variability is not the main cause of our amplitude and latency differences. Defining the exact timing of the fixation onset is important for FRP research in order to better determine latencies of the processes in question. Considering the different ways of defining this point, future studies dedicated to this issue should aim for the definition which best corresponds to unfolding of the the perceptual process. FRP research might in fact be a useful tool for determining this point in time in a phsyiololgy-relevant manner.

### N170 Adaptation

The classic N170 face effect undergoes adaptation when a target face is preceded by a face stimulus, as opposed to a stimulus from another object category (Amihai et al., 2011; Kovács et al., 2013; Nemrodov & Itier, 2011). To test for the N170 adaptation effect in free-viewing conditions, we examined sequential fixations within and between categories and their effect on the N1 amplitude. Figure 3 shows that the most extreme difference between the response to faces and eggs was obtained when the gaze shifted between categories. An important observation is that the responses to both faces and eggs in the within-category sequence were midway between the responses to faces and eggs in between-category sequences. Feuerriegel et al., 2015, found a similar effect in an ERP study and interpreted it as showing category non-specific adaptation, presumably because N1\N170 effects for both faces and chairs in their study were less negative following the presentation of a face adaptor (which we replicate with faces and eggs). However, a more plausible interpretation of both our study and theirs is that both faces and eggs elicit a unique response in the N170 time window – one more negative (faces) and one more positive (eggs, chairs) - and both show category-specific adaptation (Kovács et al., 2006). The adaptation is reflected in a category non-selective midway response when adapted in the within-category situation.

### Category Non-specific Effects

Since the contrast between faces and eggs eliminates category non-specific activity, differences between ERPs and FRPs were most prominent within each category separately (Figure 2.c, 4 and 5). The first topographic difference between viewing conditions emerged at the P1\lambda response latency, ∼100 ms from stimulus or fixation onset (Figure 4 and 5). The P1 component of ERPs is a positive deflection peaking around 100 ms after stimulus onset, maximal over occipital electrodes. It is commonly thought to reflect low-level feature processing that can be modulated by spatial attention (Hillyard & Anllo-Vento, 1998). The lambda response is a well-known component of EEG activity locked to fixation onset, with a positive peak largest at occipital sites, reversed polarity in frontal electrodes, and a peak latency ∼100 ms (Dimigen et al., 2011; Kazai & Yagi, 2005; Ries et al., 2018; Thickbroom et al., 1991; Yagi, 1981). Similar to the P1 component, the lambda response is modulated by low level features such as luminance, size and contrast (Ries et al., 2018). Based on these similarities between the P1 (in a pattern-reversal paradigm) and the lambda response, a common generator for these components has been suggested (Kazai & Yagi, 2003).

In our study, the topography of the ERP P1 component and the FRP lambda response were quite different. The FRP lambda response started earlier and was maximal in central occipital electrodes (O1/O2), where the ERP P1 was very small, if at all noticeable. The event-related P1 component was maximal in lateral occipito-temporal electrodes (P9/P10) and was of the same amplitude and latency as the fixation-related lambda response in these electrodes. These results do not necessarily contradict the possibility that there are common generators for these two components. In the study of Kazai & Yagi,, 2003, ERPs were generated by a full screen checkerboard pattern-reversal, and were compared to FRPs generated by alternate fixations on two points on the background of the same checkerboard. Presumably, the spatial extent of retinal change was similar in the two conditions. In our experiment, in contrast, the change in the retinal image in the control condition (ERPs) occurred foveally, namely, within a radius of about two visual degrees, whereas in the FRPs conditions, saccades of about eight visual degrees to the peripheral images caused an entire shift of the retinal image. The activation of a broader population of neurons responding to such a change could explain the wider distribution of the lambda response compared to the P1, requiring an additional midline dipole in our simulation. The lambda response could also be a manifestation of phase realignment of neural firing at the offset of saccades, due to the temporal predictability generated by the motor efferent copy (Purpura et al., 2003; Rajkai et al., 2008). In contrast, the timing of the stimulus onset in our Control condition was jittered, imposing an important difference in the temporal information the visual system had in this case.

During the N1 time window (∼150-200 ms), the topographies of the three conditions for each category separately were very different. In the ERP paradigm, the N1 was maximal in lateral occipito-temporal electrodes (P9/P10) but was observed in all occipital electrodes, with a positive counterpart in frontal electrodes. In contradistinction, the N1 peak of the FRPs was limited to the occipito-temporal electrodes, with no spread to the occipital electrodes, and with no positive counterpart in the frontal electrodes. As shown by the dipole simulation (Figures 6-7) this could be explained by temporal overlap of the N1 component with the strong midline occipital activity generated by an equivalent dipole with an anterior-posterior orientation (ascribed above to the lambda response). Source dipole localization on the mobile EEG data in Soto et al., 2018 resulted in one source fitted to Brodmann area 19 of the visual extrastriate cortex, accounting for the large positive lambda response, and a different source fitted to the right fusiform gyrus (Brodmann area 37). Both the dipole locations and temporal dynamics are similar to those constructed in our forward model^5^ and seem to strengthen our hypothesis that the lambda response activity overlaps the N1 time window.

A potential limitation of our experimental design is that, in the Free-viewing condition, subjects were instructed to count faces, which could confound face processing with target detection related activity. While target effects studied with ERPs are known to emerge later than 300 ms post stimulus onset (Polich, 2007), Kamienkowski et al., 2012 found evidence for early target effects (∼150 ms) unique to their Free-viewing condition, which overlap the N1 time. However, we do not believe this creates a major confound for the following reasons: 1) The early target-related effect in Kamienkowski et al’s study was found using very long fixations [550-1750 ms], and the effect was reduced when using fixations of shorter durations, as was the case in our study. 2) The P3 amplitude is known to be inversely related to target probability (Duncan-Johnson & Donchin, 1977; Polich, 2007). It is typical for an oddball condition in which targets are less frequent than non-targets while here, the probability was similar for targets and non-targets. 3) While a target response could be expected in the Free-viewing condition, this wasn’t the case for the Cued-viewing condition, in which faces and eggs had the same status. Nevertheless, we don’t see any indication for a difference between the N170 FRPs in these two conditions, suggesting that, even if any target-related response was elicited, it did not affect the N170 response we describe. 4) We examined the waveforms and topography of the response to eggs vs. faces in the Free-viewing condition. If faces were treated as targets and elicited a P3b response, we would expect parietal positivity (centered around Pz) in the comparison of faces and eggs. This, however was not the case (Supplementary Figure 6c).

## Conclusions

We demonstrate here that category specific processing can be tracked in free-viewing conditions using co-registration of eye-movements and EEG, by time-locking the activity to the first fixation on an object and disentangling activity overlap with previous and ensuing saccades. We showed that this category-specific activity is highly similar for FRPs and ERPs, provided that an appropriate contrast is used to eliminate category-non-specific response by subtraction. Nevertheless, the interpretation of FRPs to specific categories should take into account the potential interaction of the sensory or cognitive process in question with the strong evoked lambda response generated by the fixation onset. As was done over the decades for ERPs, better understanding of eye-movement related effects on the responses will allow a better interpretation of experimental effects as well as modelling of overlapping responses.

### Compliance with ethical standards: The authors declare no conflict of interests

All procedures complied with the ethical regulations of the Hebrew University of Jerusalem for experiments involving human subjects and was approved by the institutional review board and subjects signed an informed consent prior to their participation. The study was funded by a grant from the Israel Science foundation to LYD.

## Acknowledgements

This work is supported by grant 1902/14 from the Israel Science Foundation to L.Y.D. We thank Dr. Edden Gerber for his MATLAB implementation of the pre-processing procedure. We thank Chen Gueta, Namoi Revel and Chen Berkman with their help in the data collection, Oded Wertheimer for programming, Gal Chen for help with the statistical analysis and all the lab members for their valuable comments.

## Footnotes

1 In our design, categorical information of the stimulus was hindered in para-foveal view (see Materials and Methods section).

2 Saccade detection was qualitatively verified by comparison to the Engbert et al. paradigm (Engbert & Kliegl, 2003). Note that the fixations analyzed in this study follow macro-rather than micro-saccades.

3 Note we used a very large time window to preclude possible long term effects of fixation onset but this might not be necessary and adds a large number of predictors to the model.

4 Memory problems occurred when we used the default MATLAB solver for the regression analysis since our design matrix was extremely large. An alternative to solving the regression in Python is to use the LSMR solver for MATLAB as used in the Unfold toolbox (Ehinger & Dimigen, 2018)

5 Which was designed independently and prior to the publication of Soto et al.’s study.

