## Supplements for "Face Selective Neural Activity: Comparison Between Fixed and Free Viewing"

### Supplementary Material

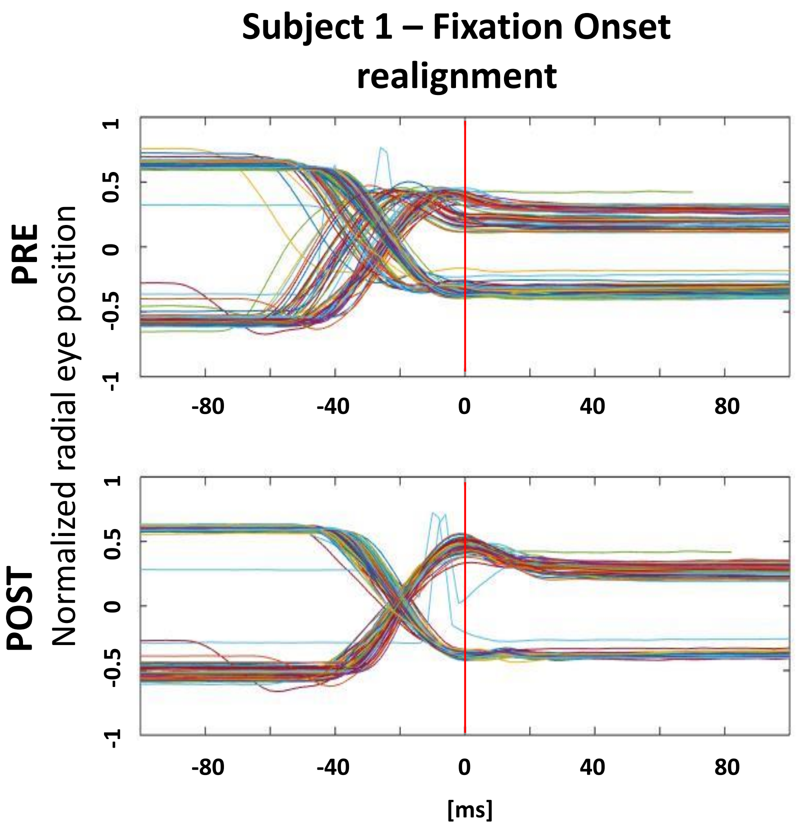


**Fig. S1** Fixation realignment results in a representative participant. Each trace depicts the eye position ±100 ms from fixation onset (red vertical line). Top: before realignment, based on the a-priori fixation onset criteria (in this case using the Eyelink default algorithm – see text). Bottom: after realignment. Fixations were realigned to the first point to cross below 2 standard deviations of the eye velocity data within this short epoch. This resulted in realigning the fixations to the end of the main saccadic movement.

**a.**

| | Cued-viewing | | Free-viewing | |
|---|---|---|---|---|
| Sub | Accuracy | pValue | Accuracy | pValue |
| 1 | 0.54 | 0.150 | 0.24 | 0.117 |
| 2 | 0.63 | 0.019 | 0.30 | 0.051 |
| 3 | 0.51 | 0.403 | 0.19 | 0.412 |
| 4 | 0.59 | 0.044 | 0.18 | 0.367 |
| 5 | 0.45 | 0.314 | 0.22 | 0.426 |
| 6 | 0.56 | 0.116 | 0.28 | 0.036 |
| 7 | 0.50 | 0.500 | 0.22 | 0.352 |
| 8 | 0.54 | 0.260 | 0.25 | 0.171 |
| 9 | 0.56 | 0.169 | 0.31 | 0.019 |
| 10 | 0.57 | 0.035 | 0.22 | 0.361 |
| 11 | 0.55 | 0.109 | 0.21 | 0.459 |
| 12 | 0.46 | 0.248 | 0.25 | 0.226 |

**b.**

| Corner Stimulus | |
|---|---|
| Sub | Accuracy |
| 1 | 0.99 |
| 2 | 0.88 |
| 3 | 0.89 |
| 4 | 0.92 |
| 5 | 0.77 |
| 6 | 0.95 |
| 7 | 0.99 |
| 8 | 0.96 |
| 9 | 0.93 |
| 10 | 0.96 |
| 11 | 0.95 |
| 12 | 0.95 |

**c.**

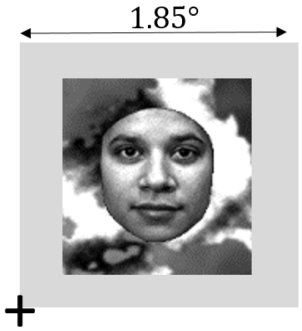


**Fig. S2** Behavioral experiment testing the ability to categorize faces and eggs (see text): a) Accuracy and p values against chance for stimuli presented in the para-foveal visual field at a distance of 9° from fixation, in the cued and free-viewing conditions. Highlighted are the subjects that preformed above chance level. Note that they are not the same subjects in both conditions. b) To test whether it is a reasonable decision to include in the analysis "First Fixations" which landed on the corner of the stimulus frame, 12 subjects were tested for their ability to categorize the faces and eggs stimuli used in the main experiment, when the corner of the stimuli are presented at fixations point (i.e., stimuli are "off-center", panel c). Subjects fixated centrally while their eyes were tracked, and stimuli were briefly presented such that one of the four corners of the stimulus appeared at the participants' fixation point in a gaze contingent manner. Stimuli appeared for 100 ms such that participants didn't have enough time to make an extra saccade. Subjects were required to press a button indicating whether the stimulus was a face or an egg. All subjects were well above chance level (all binomial $p < 10^{-15}$).

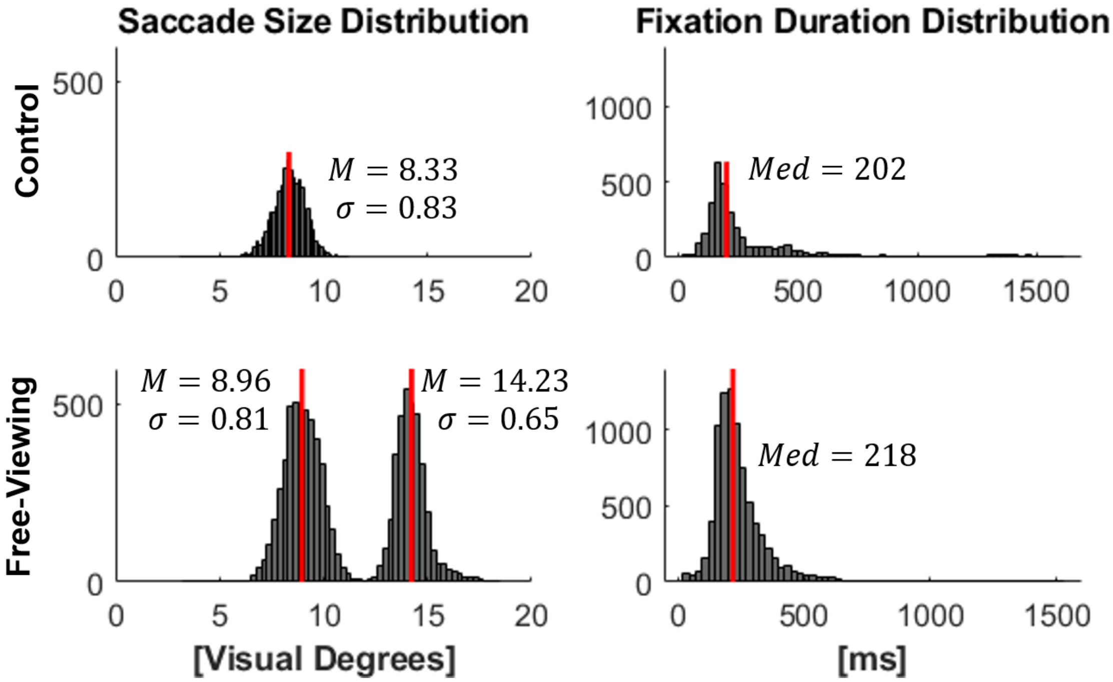


**Fig. S3 -** Distributions of saccade sizes (left) preceding the First Fixation (F1), and F1 durations (right) for the Cued-viewing (top) and Free-viewing (bottom) conditions. M – Mean, Med- Median

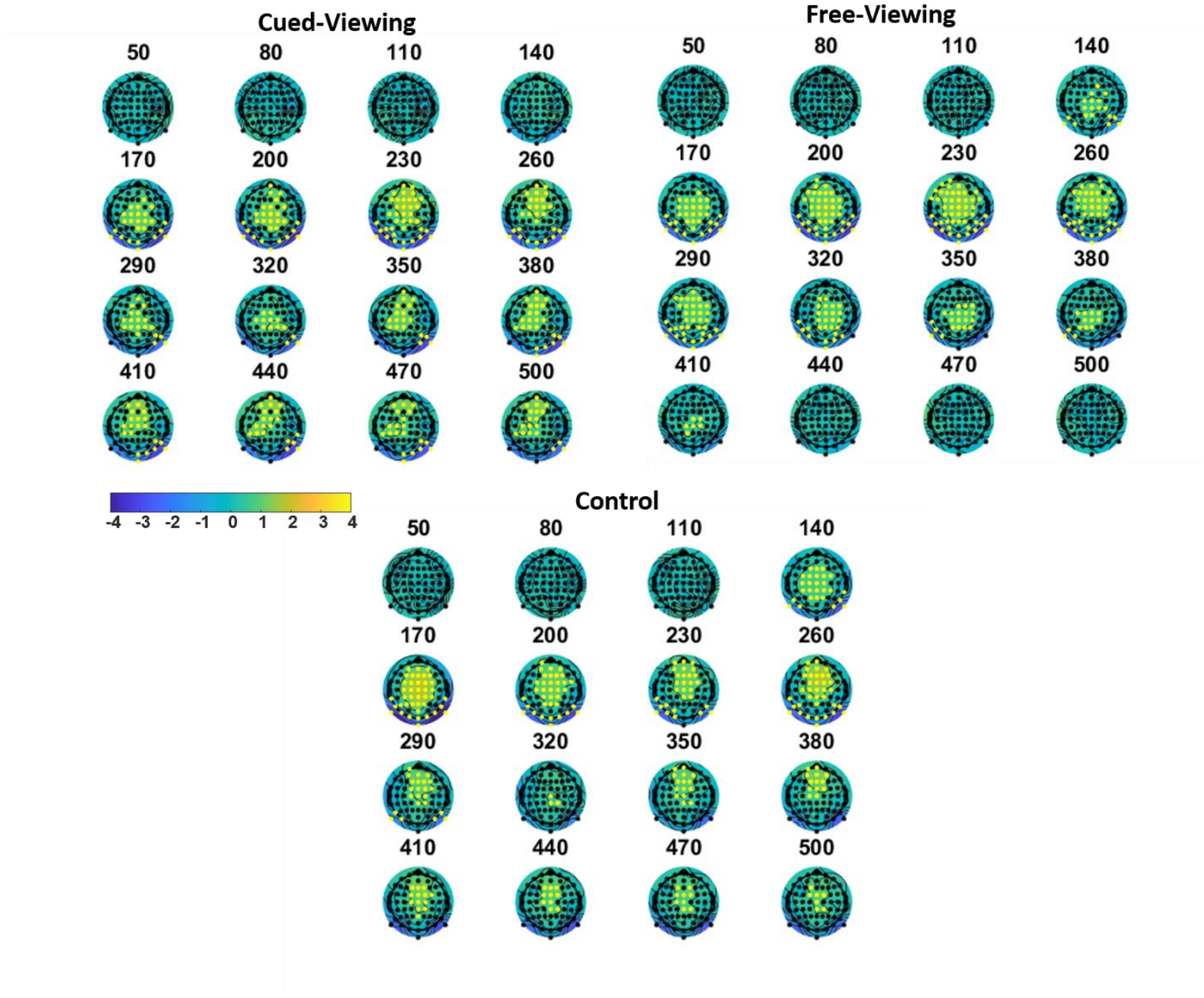


**Fig. S4** Topographies of the Face-Egg effect in each condition. The yellow dots designate electrodes which were part of a significant cluster in the 20ms window around the time above the topography. The test was done for a window of [0 800]ms but for visualization purposes we only plot the range 50-500 ms in which significant clusters were detected.

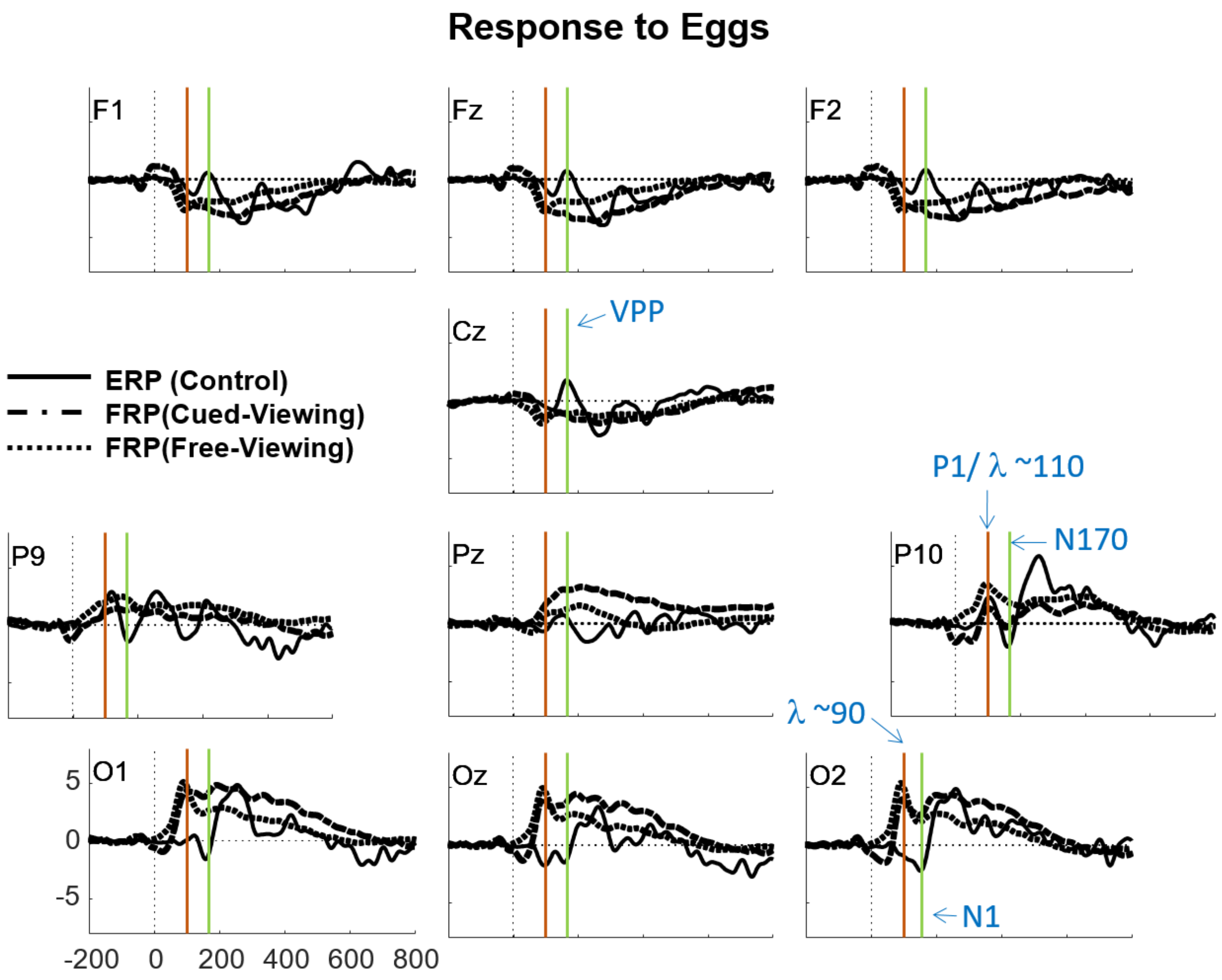


**Fig. S5** ERPs\FRPs to Eggs (as shown for faces in figure 4 of the main manuscript), depicting a selection of electrodes across the scalp.

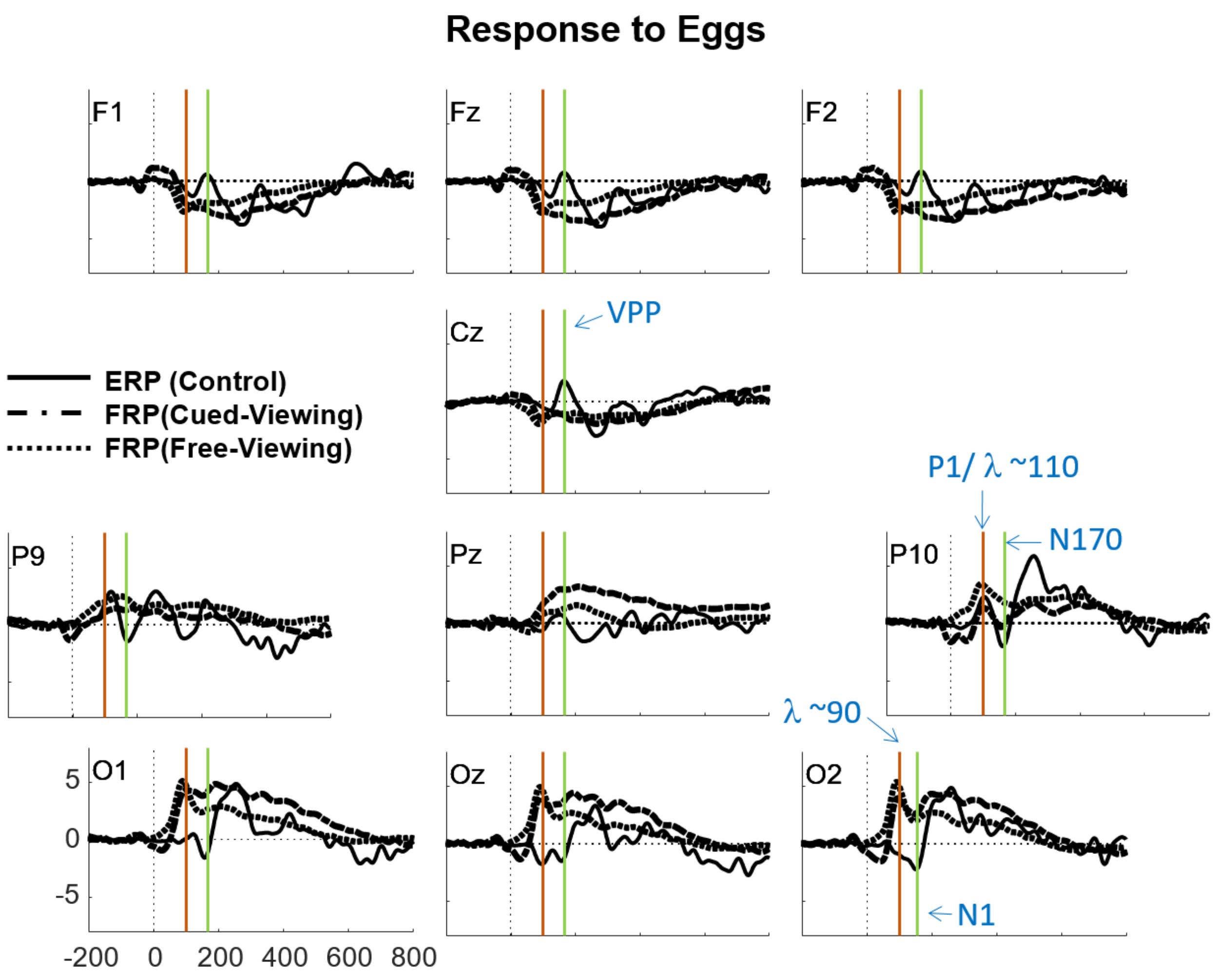

Response to Eggs
F1
Fz
F2
Cz
VPP
ERP (Control)
FRP(Cued-Viewing)
FRP(Free-Viewing)
P1/ λ ~110
N170
P9
Pz
P10
λ ~90
O1
Oz
O2
N1
5
0
-5
-200
0
200
400
600
800


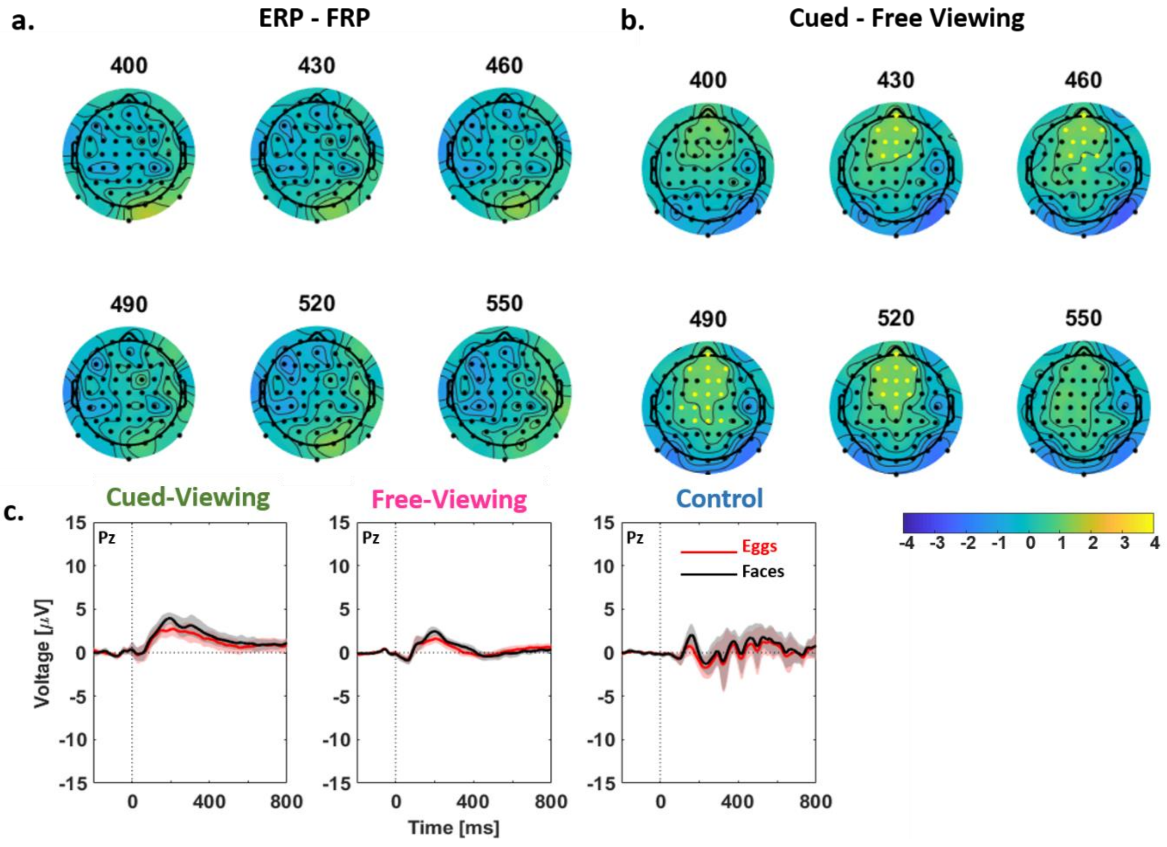

a.
ERP - FRP
400
430
460
490
520
550
b.
Cued - Free Viewing
400
430
460
490
520
550
c.
Cued-Viewing
Free-Viewing
Control
Pz
Eggs
Faces
Voltage [μV]
Time [ms]
15
10
5
0
-5
-10
-15
0
400
800
-4 -3 -2 -1 0 1 2 3 4

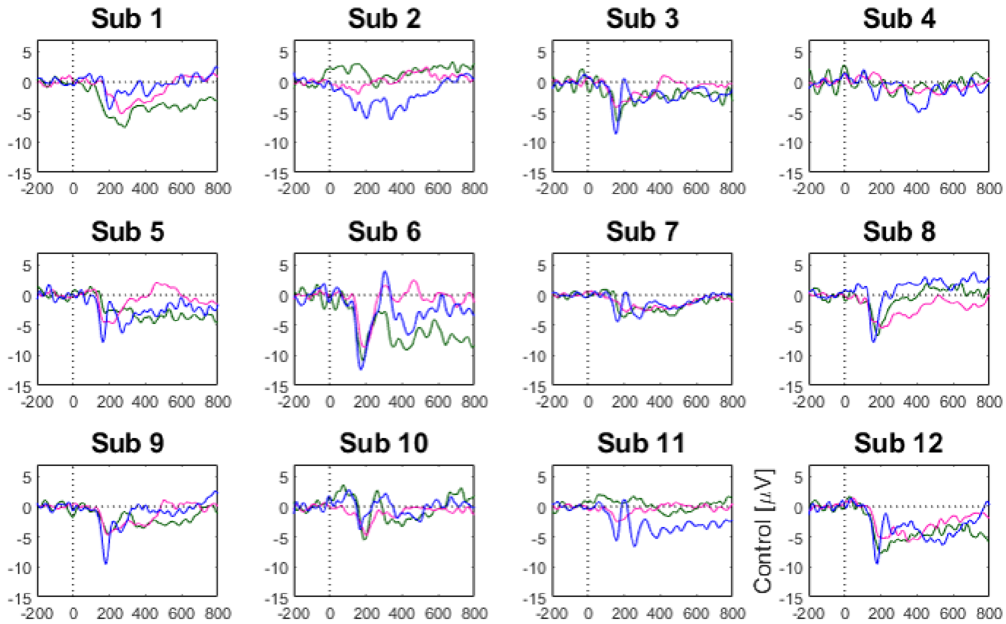


**Fig. 7 -** Single subject face-egg (face-effect) waveforms.
